# The Desmoplakin Hinge Domain Confers a Reversible Conformational Change in Response to Acute External Force

**DOI:** 10.64898/2026.10.22.760798

**Authors:** Collin M. Ainslie, William F. Dean, Yen T.B. Tran, Samuel C. Bartley, Eunjoo J. Lee, Andrew P. Kowalczyk, Daniel E. Conway, Alexa L. Mattheyses

**Affiliations:** Department of Cell, Developmental, and Integrative Biology, the University of Alabama at Birmingham, Birmingham, AL 35294, USA; Department of Biomedical Engineering, The Ohio State University, Columbus, OH 43210, USA; Departments of Dermatology and Cell and Biological Systems, College of Medicine, Pennsylvania State University, Hershey, PA 17033, USA

## Abstract

Desmosomes provide mechanical integrity to tissues that experience strain, yet how mechanical force is sensed and distributed within the desmosomal plaque remains poorly understood. Here we identify the hinge within the desmoplakin (DP) head domain is required for repositioning the DP tail in response to acute uniaxial mechanical stretch. This repositioning occurs within 30 seconds of applied stretch and reverses with release, indicating a dynamic reversible switch rather than a stable structural transition. We show this hinge-dependent repositioning correlates with the propagation of mechanical force to the desmosomal cadherins, consistent with the hinge’s role in buffering tension at the cadherin interface during mechanical loading. Deletion or shortening of the hinge abolishes the stretch-induced response and compromises desmosome mechanical integrity. The repositioning of the DP tail in the calcium-dependent to hyper-adhesive transition was not impacted by alterations to the hinge, demonstrating that DP repositioning due to acute force or adhesive state is mechanistically distinct. Together, these findings define a reversible, hinge-dependent mechanotransduction pathway through which DP senses and relays mechanical force within the desmosome. This mechanism could act as a mechanical buffer by reducing the impulse of forces on the cadherins, and play a central role in mechanical resilience.

## Introduction

The outermost layer of the skin is a stratified epithelium that provides the body’s first line of defense against the outside environment. Keratinocytes, its most abundant cell type, are held together by adhesive cell-cell junctions including desmosomes^1^. A major function of desmosomes is to maintain tissue integrity by resisting mechanical forces. To effectively resist mechanical forces, desmosomal proteins must work together harmoniously. The desmosomal cadherins desmogleins (Dsgs) and desmocollins confer adhesion though *trans*-binding. Their cytoplasmic tails bind the armadillo proteins, plakophilin and plakoglobin which in turn bind the cytoskeletal linker protein desmoplakin (DP). DP anchors the desmosome complex to intermediate filaments, such as keratin in skin^2^ (Fig. 1a). Disruption of any of these proteins is associated with a variety of disorders, many of which share a phenotype of weakened desmosomal adhesion^3,4^. How these proteins work together to confer the mechanical properties of desmosomes has yet to be established.

**Figure 1:**
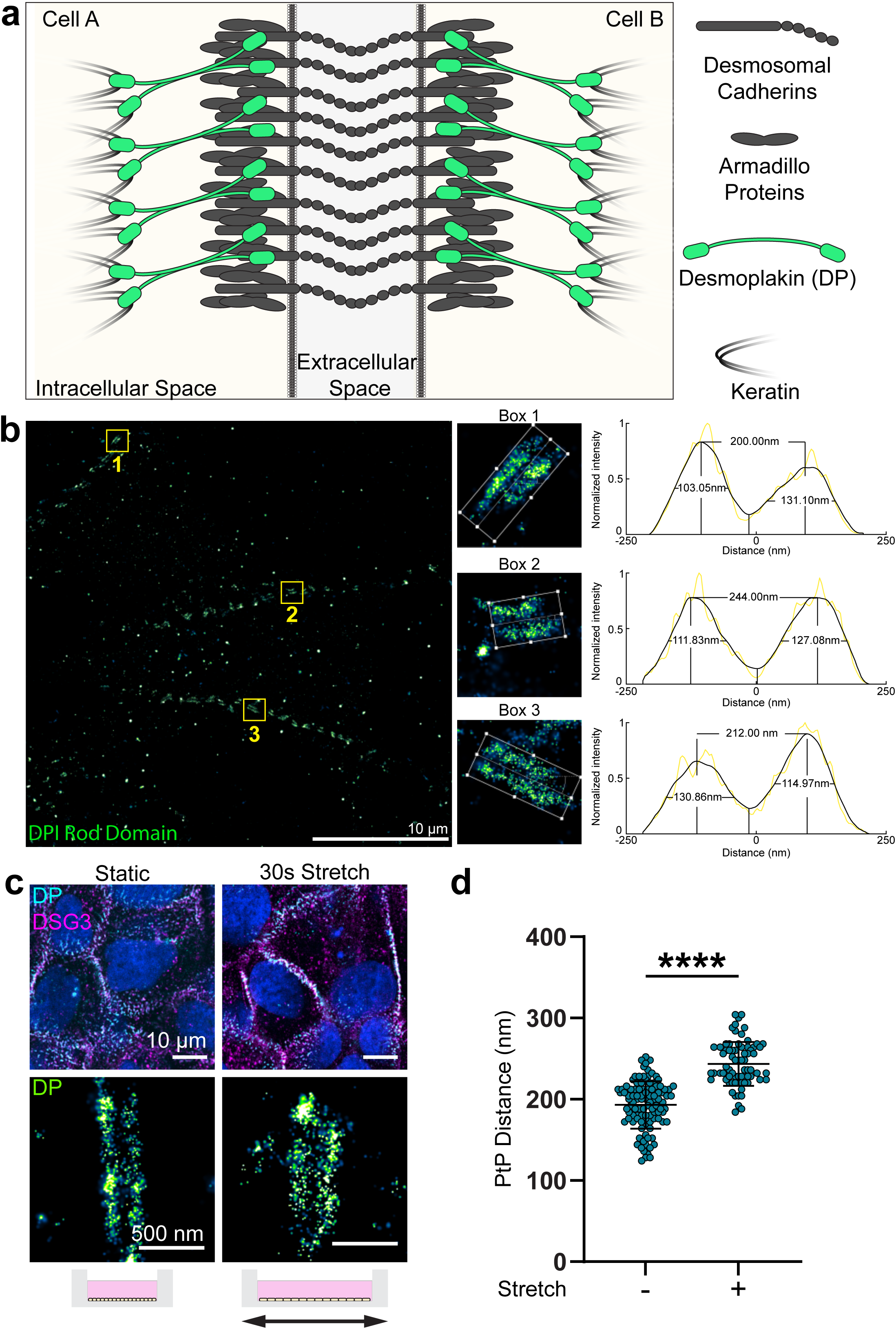
DP is architecturally flexible in desmosomes. **a**, Schematic of an individual desmosome spanning two neighboring cells with desmosomal cadherins, armadillo proteins, desmoplakin, and keratin. **b,** Representative dSTORM image of the DPI rod domain in WT HaCaT cells in 1-day confluent culture on glass coverslips. The ROI zoom-ins show the length and orientation of each desmosome as defined for analysis, with plots of intensity as a function of position across each desmosome. The distance between the peaks was quantified as the plaque-to-plaque (PtP) distance. Scale bar = 10 μm **c,** Representative confocal and dSTORM images of DPI in static and stretched WT HaCaT cells. The DPI rod domain was labeled with Alexa Fluor 488 in confocal images and Alexa Fluor 647 in dSTORM images, Dsg 3 is labeled with Alexa Fluor 647 in confocal images. Confocal scale bar = 10 μm dSTORM scale bar = 500 nm **d,** Scatter plot of the DPI rod domain PtP distance from static and stretched WT HaCaT cells (Static: n = 103, 193 ± 29 nm; Stretched: n = 73, 244 ± 27 nm. n = number of desmosomes; mean ± sd). Desmosomes were analyzed from three independent replicates. Shapiro-Wilks test assessed normality, and PtP distance was compared with a Mann-Whitney U test (****p < 0.0001).

Dsg2, Dsg3, and DP have all been shown to be mechanically sensitive^5–8^. While cadherin tension may be isoform and context-dependent, DP is not under tension in static conditions and is loaded upon external mechanical forces or actomyosin contractility^5–8^. This introduces an interesting paradigm of tension across the desmosome complex, suggesting the desmosomal cadherins in the core can be under load while DP only experiences load under mechanical forces.

DP is mechanically sensitive, and while it is not under tension in static conditions, upon experiencing mechanical forces it becomes loaded and responds architecturally through tail domain repositioning^2,9–11^. When DP experiences mechanical forces, the C-terminal tail domain repositions away from the plasma membrane while the N-terminal head domain position does not change. This change has been observed in cells with high actomyosin contractility, cardiomyocytes, and cells under external strain^7,8^. The mechanism behind this rearrangement remains unknown, with a conformational change of the DP head domain as a leading hypothesis^7,8^. The DP head domain has an unstructured N-terminal region followed by four spectrin repeat (SR) domains (SR3-SR6), with an SH3 domain between SR4-SR5 constituting the long arm and SR7-8 comprising the short arm. A 37 amino acid unstructured hinge sequence connects the two arms^10,12,13^. The head domain can adopt elongated “I” and folded “U” conformations, proposed to be made accessible by the flexible hinge between the two arms^10^. It is hypothesized that the DP head domain undergoes a conformational change at the hinge in response to applied mechanical force, leading to a repositioning of the tail domain^7,8^.

Here we developed a novel series of DP head domain structural variants which, combined with super-resolution direct stochastic optical reconstruction microscopy (dSTORM), revealed the hinge is essential for the DP architectural response to mechanical stretch. We established DP’s response is both rapid and reversible: within 30 s of applied stretch, the head domain opens at the hinge and then closes following 30 s of release of stretch. Additionally, with the opening and closing of the DP head domain, we measured tension at Dsg3. From this, we define a reversible mechanotransduction cascade arising from stretch leading to DP opening and tension on the cadherins. This suggests DP’s response to mechanical forces could serve as a buffer to dampen force distribution throughout the desmosomal complex, shielding the cadherins. These findings provide novel insights into DP as a mechanosensitive molecule, as well as force propagation through the desmosome in epithelial tissue.

## Results

### DP Tail Domain Architecture Changes with Acute Mechanical Force

Desmosome ultrastructure consists of the extracellular domain (ECD) where trans-binding occurs between desmosomal cadherins, the outer dense plaque (ODP) containing the cytoplasmic tails of the desmosomal cadherins, the armadillo proteins, and the DP head domain, and the inner dense plaque (IDP) containing the DP tail domain while the DP rod domain spans the ODP and IDP^14,15^ (Fig. 1a). This overall protein architecture is variable with changes in protein isoforms and localization of the DP tail domain based on cell location within a tissue, desmosome maturation, and mechanical force ^7,8,16,17^. Changes in the DP tail domain position indicate that desmosome architecture is dynamic, but the mechanism underlying this remains unclear. To address this, we use super-resolution dSTORM to resolve single desmosomes as two parallel DP plaques (Fig. 1b). The DP architecture is characterized by the plaque-to-plaque (PtP) distance, quantified as the distance between peaks obtained from a series of intensity linescans perpendicular to each desmosome midline^17^.

To examine how DP architecture responds to acute mechanical force, wild type (WT) HaCaT cells were grown on a PDMS membrane, subjected to 30% uniaxial stretch for 30 s, fixed in the stretched position, and labeled with an antibody to the C-terminal end of the DPI rod domain (amino acid 1900 – 1950). Cells cultured on the same substrate but not subjected to stretch served as static controls (Fig. 1c). We found a significant increase in DP PtP distance in cells subjected to stretch compared to the static control (static, 192 ± 25 nm; stretch, 242 ± 27 nm). This rapid response to acute stretch agrees with independent observations that DP repositions in cells with high actin contractility or after a prolonged 30 m stretch, indicating this switch is not specific to one mode or duration of force application^7,8^. These findings suggest a general mechanosensing property of DP that allows for its architectural repositioning to mechanical forces. We next sought to define the mechanism underlying this observation.

### DP Head Domain Variants

One hypothesis is that the head domain transitions from a no/low stress folded state to an open, more linear state under tension^7,8,10^. To determine whether the DP head domain mechanically mediates tail domain position, we generated three structural variants of full-length WT DPI with a C-terminal mEGFP, each containing deletions or substitutions in the head domain (Fig. 2a). The hinge (AA 627-663) was deleted to generate ΔHinge or replaced with the shorter 7 amino acid hinge from plakin family member envoplakin to generate Hinge>ENVP. To control for contributions of nearby domains, the SR6 domain was deleted to generate ΔSR6. WT DPI and each variant were stably expressed in DP KO HaCaT cells ^8,10^.

**Figure 2:**
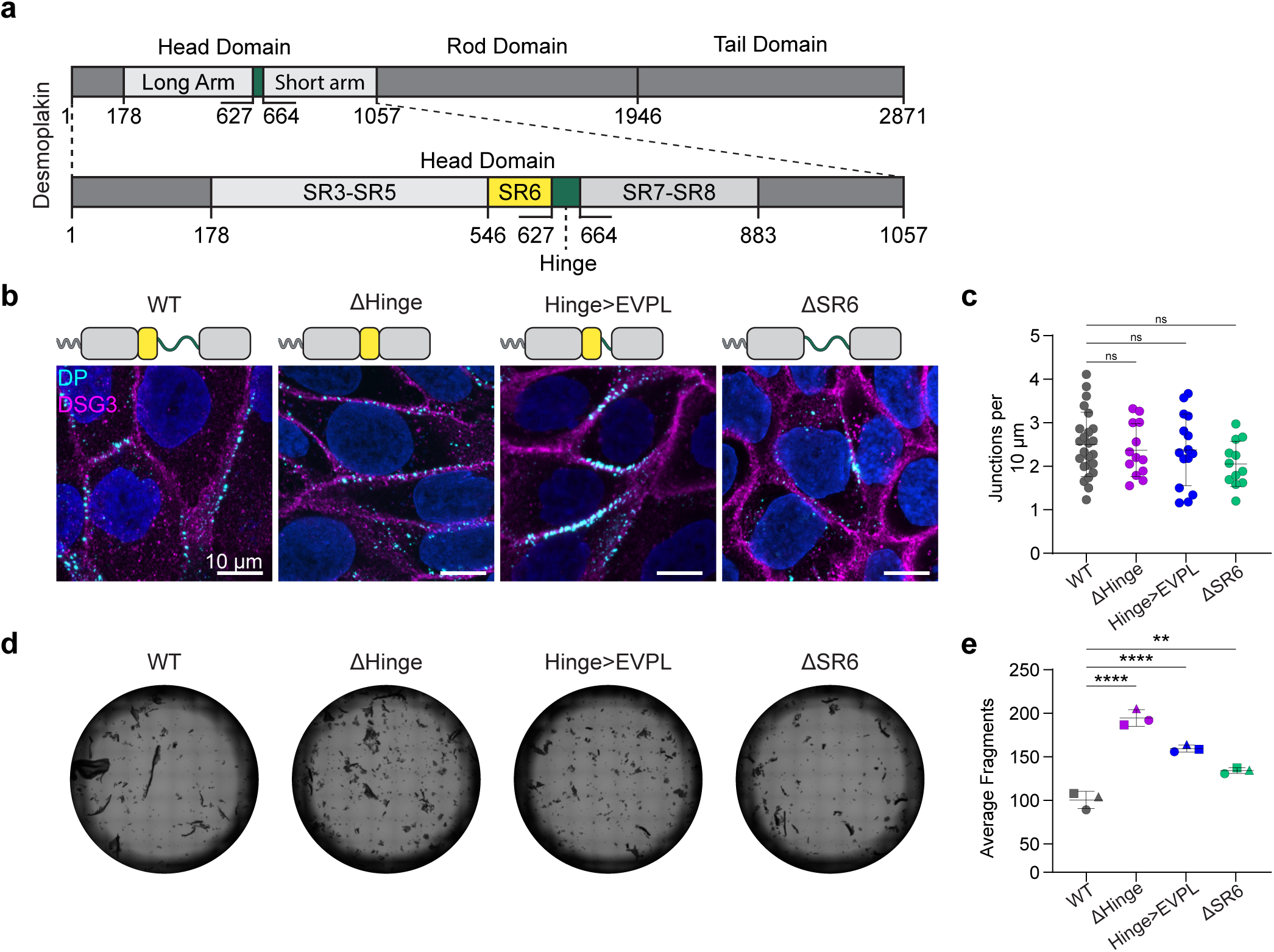
DP head domain structural variants. **a,** schematic representation of DP AA sequence highlighting the head domain regions. **b,** Representative confocal images of DP KO HaCaT cells stably expressing DP WT and structural variants with an anti-GFP nanobody conjugated to Atto 488 and Dsg 3 labeled with Alexa Fluor 647. Scale bar = 10 μm. Graphical representations of DP structural variants (WT, ΔHinge, Hinge>EVPL, and ΔSR6). **c,** Scatter plot of the number of desmosomes per 10 μm for the DPI variant stable cells (WT: n = 24, 2.5 ± 0.7; ΔHinge: n = 13, 2.4 ± 0.6; Hinge>EVPL: n = 15, 2.4 ± 0.8; ΔSR6: n = 12, 2.0 ± 0.5. n = number of cells; mean ± sd). **d,** Representative brightfield images of the dispase fragmentation assay. **e,** Scatter plot of the average number of fragments in the dispase fragmentation assay (WT: 101 ± 10; ΔHinge: 195 ± 10; Hinge>EVPL: 160 ± 4; ΔSR6: 134 ± 3. n = 3 biological replicates, each with 3 technical replicates; mean ± sd). A Shapiro-Wilks confirmed normality (p>0.05). An ordinary one-way ANOVA and Dunnett’s multiple comparisons test assessed desmosomes per 10 μm (ns not significant). Ordinary one-way ANOVA with Dunnett’s multiple comparison test assessed average dispase fragments. (**** p<0.0001, ** p=0.0013, ns not significant).

Confocal imaging confirmed the DP structural variants co-localize with Dsg3 in puncta at cell-cell borders (Fig. 2b). Quantitative analysis revealed no significant differences in junction abundance compared to WT, suggesting the DP structural variants do not interfere with desmosome assembly (Fig. 2c). Desmosome function was tested with a dispase fragmentation assay, which revealed more fragmentation and weaker adhesion in the structural variants compared to WT. ΔHinge provided the least protection from mechanical forces while Hinge>ENVP and ΔSR6 were intermediate (number of fragments: WT, 101 ± 10; ΔHinge, 195 ± 10; Hinge>ENVP, 160 ± 4; ΔSR6 134 ± 3) (Fig. 2d,e). Together these data suggest the DP structural variants incorporate into desmosomes but weaken cell-cell adhesion.

Next, we asked whether the DP head domain contributes to the architectural remodeling observed in desmosome maturation. As desmosomes mature in culture, the DP tail domain moves closer to the plasma membrane (decreased PtP distance), a shift also seen in hyper-adhesive desmosomes generated by plakophilin 1 overexpression^16,17^. This maturation-associated repositioning reflects the difference seen between less mature basal and more mature supra-basal keratinocytes in human epithelial tissue^17^. To establish a straightforward experimental paradigm to examine the correlation between DP architecture and desmosome maturity, WT HaCaT cells were grown to 1-day or 5-day confluence to mimic less mature calcium-dependent and more mature hyper-adhesive junctions (Fig. 3a). dSTORM revealed a significant 66 ± 40 nm decrease in the PtP distance of DPI in cells in a 5-day confluent culture compared to cells in a 1-day confluent culture (5-day, 152 ± 27 nm; 1-day, 218 ± 30 nm) (Fig. 3a,b). To determine whether the DPI structural variants share this repositioning phenotype, the stably expressing HaCaT cells were grown to 1-day or 5-day confluence (Fig. 3c,d). The PtP distance in WT expressing cells decreased by 55 ± 45 nm in 5-day confluent culture compared to 1-day confluent culture, similar to what was measured in WT HaCaT cells (1-day, 231 ± 33 nm; 5-day, 176 ± 30 nm). The PtP distance decreased in 5-day confluent culture when compared to 1-day confluent culture for ΔHinge: 64 ± 50 nm, DPI-EVPL: 62 ± 51 nm, and DPI-ΔSR6: 54 ± 48 nm. (1-day/5-day: ΔHinge, 256 ± 37 nm / 192 ± 34 nm; Hinge>EVPL, 240 ± 36 nm / 178 ± 36 nm; ΔSR6, 252 ± 38 nm / 198 ± 29 nm) (Fig. 3e). We also noted the PtP distance in 1-day confluent culture was not the same for all variants, with ΔHinge and ΔSR6 having significantly larger PtP distances than WT. This suggests these domains may change the overall shape of the protein, but do not limit the possible orientations of the tail. Regardless of these differences, the structural variants all underwent a similar repositioning of the DP tail domain from 1-day to 5-day culture, validating their ability to respond architecturally to maturation cues similar to WT DP. These findings suggest the mechanism underlying DP tail domain repositioning toward the plasma membrane in maturation is independent of the hinge region.

**Figure 3:**
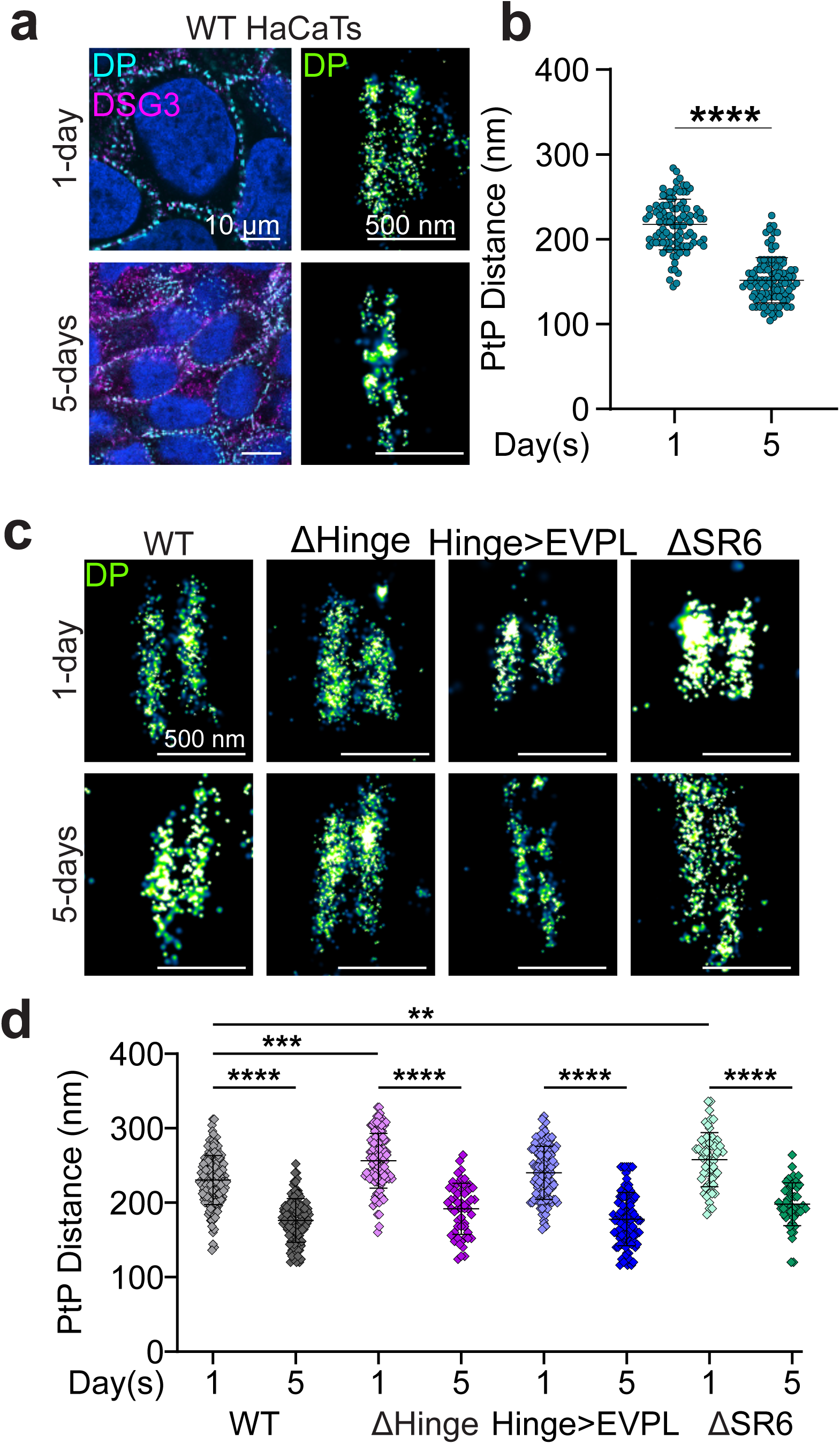
DP repositioning in maturation is not dependent on the hinge. **a,** Representative confocal and dSTORM images of DPI in 1- and 5-day confluent junctions in WT HaCaTs. The DPI rod domain was labeled with Alexa Fluor 488 in confocal and Alexa Fluor 647 in dSTORM images, Dsg 3 is labeled with Alexa Fluor 647 in confocal images. Confocal scale bar = 10 μm, dSTORM scale bar = 500 nm **b,** Scatter plot of DPI rod domain plaque-to-plaque (PtP) distances in 1-day or 5-day confluent WT HaCaTs (1 day: n = 100, 218 ± 30 nm; 5-day: n =106, 152 ± 27 nm. n = number of desmosomes; mean ± sd). **c,** Representative dSTORM images of DPI-mEGFP constructs in 1-day confluent or 5-day confluent culture, labeled with an anti-GFP nanobody conjugated to Alexa Fluor 647. scale bar = 500 nm **e,** Scatter plot of the PtP distance of the tail domain of the DPI-mEGFP constructs in a 1-day or 5-day confluent culture (WT 1-day confluent: n = 162, 231 ± 33 nm; WT 5-day confluent: n = 108, 176 ± 30 nm; ΔHinge 1-day confluent: n = 108, 256 ± 37 nm; ΔHinge 5-day confluent: n = 55, 192 ± 34 nm; Hinge>EVPL 1-day confluent: n = 110, 240 ± 36 nm; Hinge>EVPL 5-day confluent: n = 97, 178 ± 36 nm; ΔSR6 1-day confluent: n = 66, 252 ± 36 nm; ΔSR6 5-day confluent: n = 48, 198 ± 29. n = number of desmosomes; mean ± sd). Shapiro-Wilks test assessed for normality. Mann-Whitney U test assessed the PtP distances of WT HaCaT cells (**** p<0.0001). Kruskal-Wallis with Dunn’s multiple comparisons test assessed the significant differences in PtP distance of DPI head domain constructs (****p<0.0001, *** p<0.001, ** p<0.01).

### The DP Hinge is Mechanosensitive

Next, we asked whether the force-dependent repositioning observed for the tail domain could be explained by a tension-induced conformational change in the DP head domain (Fig. 1c,d). To address this, we subjected HaCaT cells stably expressing the DP structural variants to 30% uniaxial stretch for 30 s. Confocal imaging confirmed membrane localization of the DP variants in both static and stretched cells (Fig. 4a). dSTORM revealed that following stretch, the WT PtP distance increased by 30 ± 62 nm (static, 220 ± 39 nm; stretch, 250 ± 48 nm), indicating a repositioning of the tail domain similar to that measured in WT HaCaTs (Fig. 4b,c). In contrast, when the hinge domain was deleted or shortened, there was no significant change in PtP distance following mechanical stretch (static / stretch: ΔHinge, 257 ± 38 nm / 253 ± 43 nm; Hinge>ENPL 257 ± 39 nm / 255 ± 43 nm) (Fig. 4b,c). Deletion of the SR6 domain led to a 46 ± 70 nm increase in tail domain PtP distance in response to mechanical stretch (static, 234 ± 49 nm; stretch, 280 ± 50 nm) (Fig. 4b,c). The DP N-terminus PtP distance was consistent in static and stretched junctions (static, 83 ± 10 nm; stretch, 80 ± 12 nm), suggesting the observed changes cannot be explained by an overall movement of the entire DP molecule away from the membrane and does not underlie the observed changes in tail domain localization (Fig. 4d). Instead, the absence of a measured change in PtP distance only for variants with a deleted or shortened hinge region suggests the DP head is mechanically sensitive; with the flexible hinge serving as the primary driver for architectural rearrangement in response to applied force.

**Figure 4:**
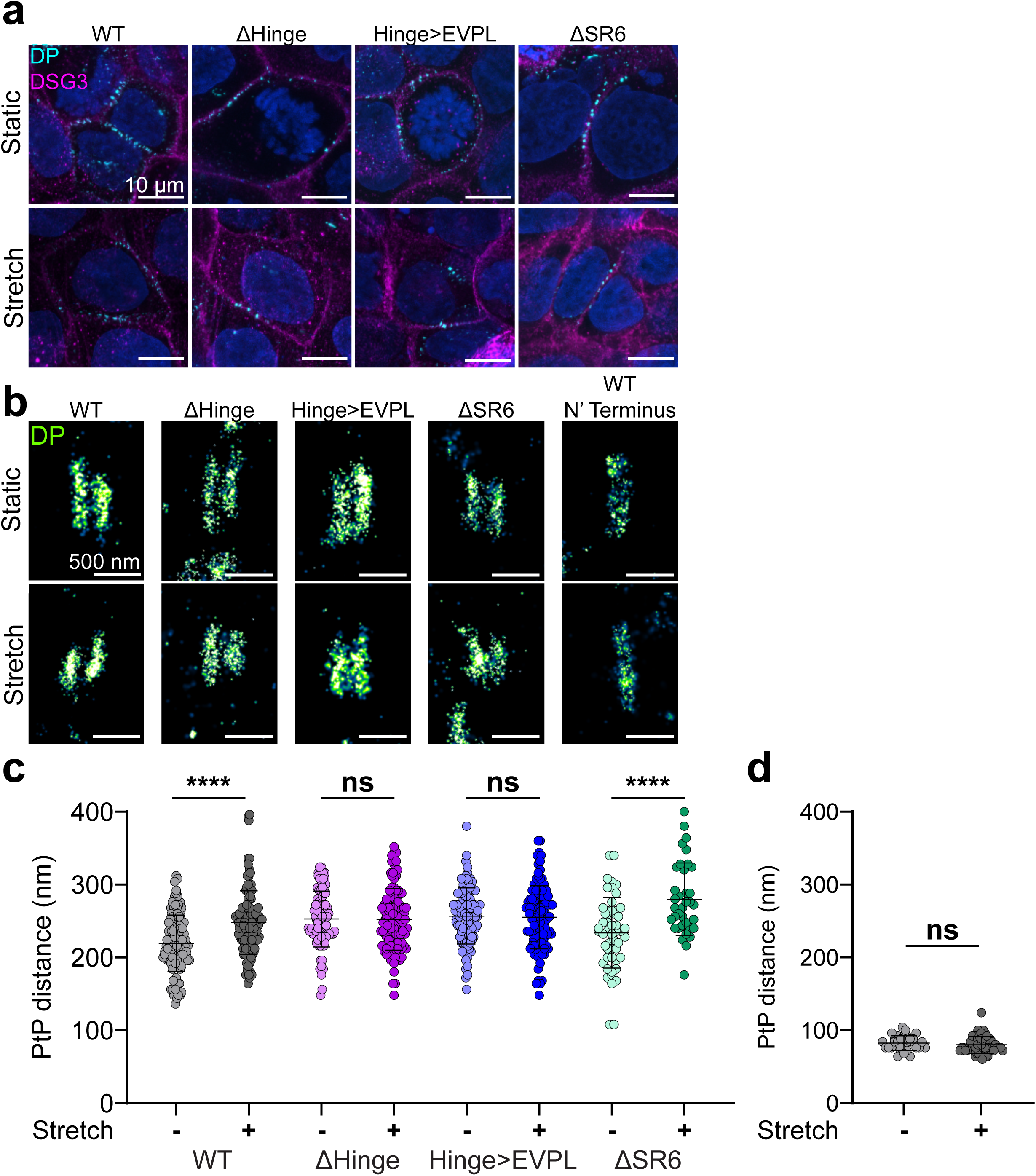
The DP Hinge is Essential for Tail Domain Relocation in Response to Stretch. **a**, Representative confocal images of static or 30 s stretched WT, ΔHinge, Hinge>EVPL, and ΔSR6 labeled with an anti-GFP nanobody conjugated to Atto 488 and Dsg 3 labeled with Alexa Fluor 647. scale bar = 10 μm **b,** Representative dSTORM images of desmosomes from static or stretched cells WT, ΔHinge, Hinge>EVPL, ΔSR6, and WT N-terminus. scale bar = 500 nm **c,** Scatter plot of the tail domain plaque-to-plaque (PtP) distance of DPI-mEGFP constructs in static and stretch conditions (WT static: n = 115, 220 ± 39 nm; WT stretch: n = 113, 248 ± 43 nm; ΔHinge static: n = 43, 258 ± 38 nm; ΔHinge stretch: n = 60, 253 ± 43 nm; Hinge>EVPL static: n = 100, 257 ± 39 nm; Hinge>EVPL stretched: n =107, 255±43 nm; ΔSR6 static: n = 53, 234 ± 49 nm; ΔSR6 stretched: n = 36, 280 ± 50 nm. n = number of desmosomes; mean ± sd). **d,** Scatter plot of the N-terminus PtP distance of WT in static and stretched junctions (WT N-terminus static: n = 31, 83 ± 10 nm; WT N-terminus stretch: n = 52, 80 ± 12 nm. n = number of desmosomes; mean ± sd). Shapiro-Wilks test assessed for normality. Mann-Whitney U test assessed the significant differences of PtP distances of WT (**** p<0.0001). Welch’s t-test assessed the significant differences of PtP distances of ΔHinge, Hinge>EVPL, and ΔSR6 (**** p<0.0001, ns not significant) Mann-Whitney U test assessed significant difference of WT N-terminus (ns not significant).

### DP responds to force direction and duration of force

We next examined if the mechanical response of DP depends on desmosome orientation relative to the direction of applied mechanical force. Desmosomes were classified based on their orientation relative to the stretch axis, where 0° represents a desmosome with its plaques oriented parallel to stretch axis and 90° represents desmosomes oriented perpendicular to the stretch axis (Fig. 5a). Desmosomes oriented perpendicular to the stretch axis had larger PtP distances compared to those parallel to the stretch axis (0°-30°, 229 ± 42 nm; 30°-60°, 251 ± 35 nm; 60°-90°, 273 ± 60 nm) (Fig. 5b). As anticipated, there was no statistical difference in PtP distance based on orientation to stretch axis in control unstretched cells (0°-30°, 226 ± 39 nm; 30°-60°, 218 ± 43 nm; 60°-90°, 214 ± 32 nm) (Fig. 5b). Desmosomes expressing the DP structural variants revealed no orientation-dependent differences in PtP distances relative to the stretch axis (Fig. S1). This suggests desmosome orientation relative to mechanical force correlates with the extent of its architectural response, and that orientation did not mask architectural changes in our DP head domain variant constructs.

**Figure 5:**
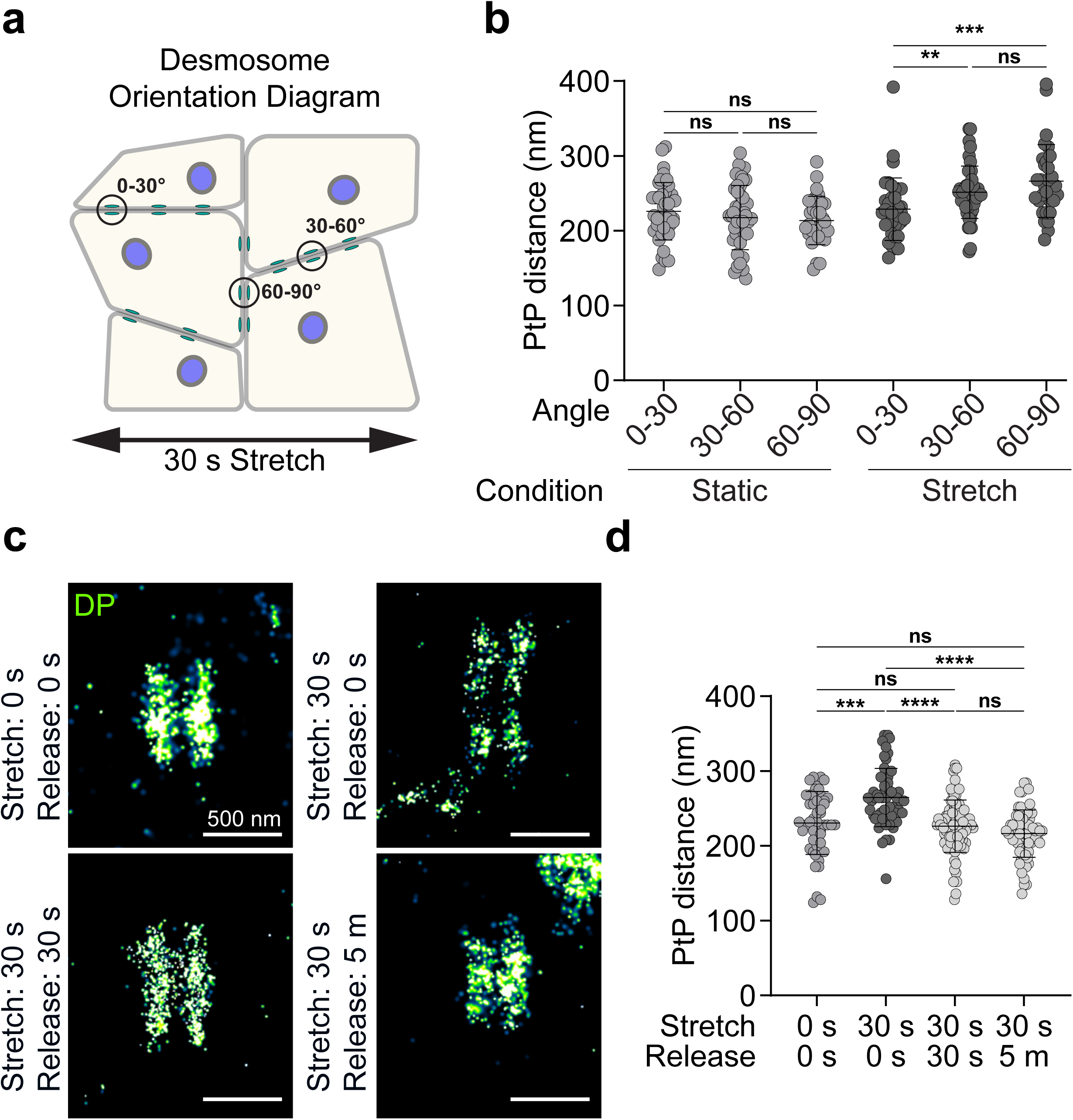
The DP Head Domain Opens and Closes Rapidly in Response to Mechanical Forces. **a**, Diagram illustrating desmosome orientation to mechanical stretch. **b,** Scatter plot of the plaque-to-plaque (PtP) distances of WT in static or stretched junctions sorted by angle relative to stretch (Static 0-30: n = 41, 226 ± 39 nm; Static 30-60: n = 43, 218 ± 43 nm; Static 60-90: n = 31, 214 ± 32 nm; Stretch: 0-30: n = 37, 229 ± 42 nm; Stretch 30-60: n = 46, 251 ± 35 nm; Stretch 60-90: n = 30, 266 ± 49 nm. n = number of desmosomes; mean ± sd). **c,** Representative dSTORM images of static, 30 s stretched no release, 30 s stretch with 30 s release, and 30 s stretch with 5 m release of WT labeled with an anti-GFP nanobody conjugated to Alexa Fluor 647. scale bar = 500 nm **d,** Scatter plot of the PtP distances of the tail domain of static, 30 s stretched no release, 30 s stretched with 30 s release, and 30 s stretch with 5 m release WT (static: n = 48, 231 ± 42 nm; 30 s stretched no release: n = 54, 265 ± 39 nm; 30 s stretch and 30 s release: n = 90, 226 ± 35 nm; 30 s stretch and 5 m release: n = 69, 217 ± 32 nm. n = number of desmosomes; mean ± sd). Shapiro-Wilks test assessed for normality. One-way ANOVA followed by Dunnett’s multiple comparisons test revealed no significant differences in PtP of static junctions sorted by orientation to the stretch axis (ns not significant). Kruskal-Wallis with Dunn’s multiple comparison test assessed significant differences in PtP distances of stretched junctions sorted by orientation to the stretch axis (*** p<0.001, ** p<0.01, ns not significant). Kruskal-Wallis with Dunn’s multiple comparison test assessed significant differences in PtP distances of junctions released from stretch (**** p<0.0001, *** p<0.001, ns not significant).

When DP experiences mechanical force, it opens. Next, we wanted to explore the time domain and test if DP closes following release of force. WT DP expressing cells were subjected to 30% uniaxial stretch for 30 s followed by release of stretch for 30 s or 5 m (Fig. 5c). dSTORM imaging and analysis revealed the DP tail domain PtP distance returned to the pre-stretch values 30 s after release of stretch (static, 225 ± 48 nm; 30 s stretch, 255 ± 40 nm; 30 s release, 236 ± 32 nm; 5 m release, 219 ± 31 nm) (Fig. 5d). This is consistent with a rapid and reversible opening and closing of the head domain in response to both loading and unloading, similar to the dynamics of tension on DP previously reported^10^.

### DP Head Domain Opening is Part of a Mechanical Cascade

To understand how force propagates through the desmosome to the adhesive core, we introduced a FRET based Dsg 3 tension sensor (Dsg3-TS) to the stretch experiments^6^. For simultaneous imaging of DP localization and Dsg 3 tension, we generated a construct containing full-length DPI with a C-terminal halo tag and stably expressed it in DP KO HaCaT cells. We found WT DPI-halo localized with Dsg 3 on cell borders, similar to WT DPI-mEGFP (Fig. S2). We then transfected Dsg3-TS into the DPI-Halo stable cells grown on a flexible PDMS membrane and subjected to 30 s or 10 m of 30% uniaxial stretch. Quantification of the FRET ratio revealed a significant increase in tension across Dsg 3 in cells stretched for 30 s and 10 m compared to static controls. In cells stretched for 30 s and then released from stretch for 30 s, the Dsg 3 tension was lost and returned to baseline (static, 0.71 ± 0.15; 30 s stretch, 0.55 ± 0.07; 10 m; stretch, 0.6 ± 0.09; 30 s release, 0.72 ± 0.11). There was no change in FRET ratio in DPI-Halo cells expressing the Dsg3 tailless control upon experiencing mechanical stretch (Fig. S3). This suggests that upon application of acute force, tension is reversibly propagated to Dsg 3 within the same time frame as the DP head domain conformational change.

## Discussion

In this work, we establish that the DP head domain is mechanically sensitive and required for DP tail movement in response to acute mechanical force. Desmosomes respond to force with a proposed opening of the DP head domain, leading to a repositioning of the tail, and a force across the complex is felt by DP^18^ and cadherins. This process is reversible, and the release of force allows refolding of the head domain and loss of tension on DP^18^ and Dsg 3, thereby resetting the desmosome back to baseline (Fig. 7). Deletion or shortening of the DP hinge abrogated the force induced-repositioning of the tail domain in response to stretch, indicating the hinge plays a central role in the architectural response to applied force.

**Figure 6:**
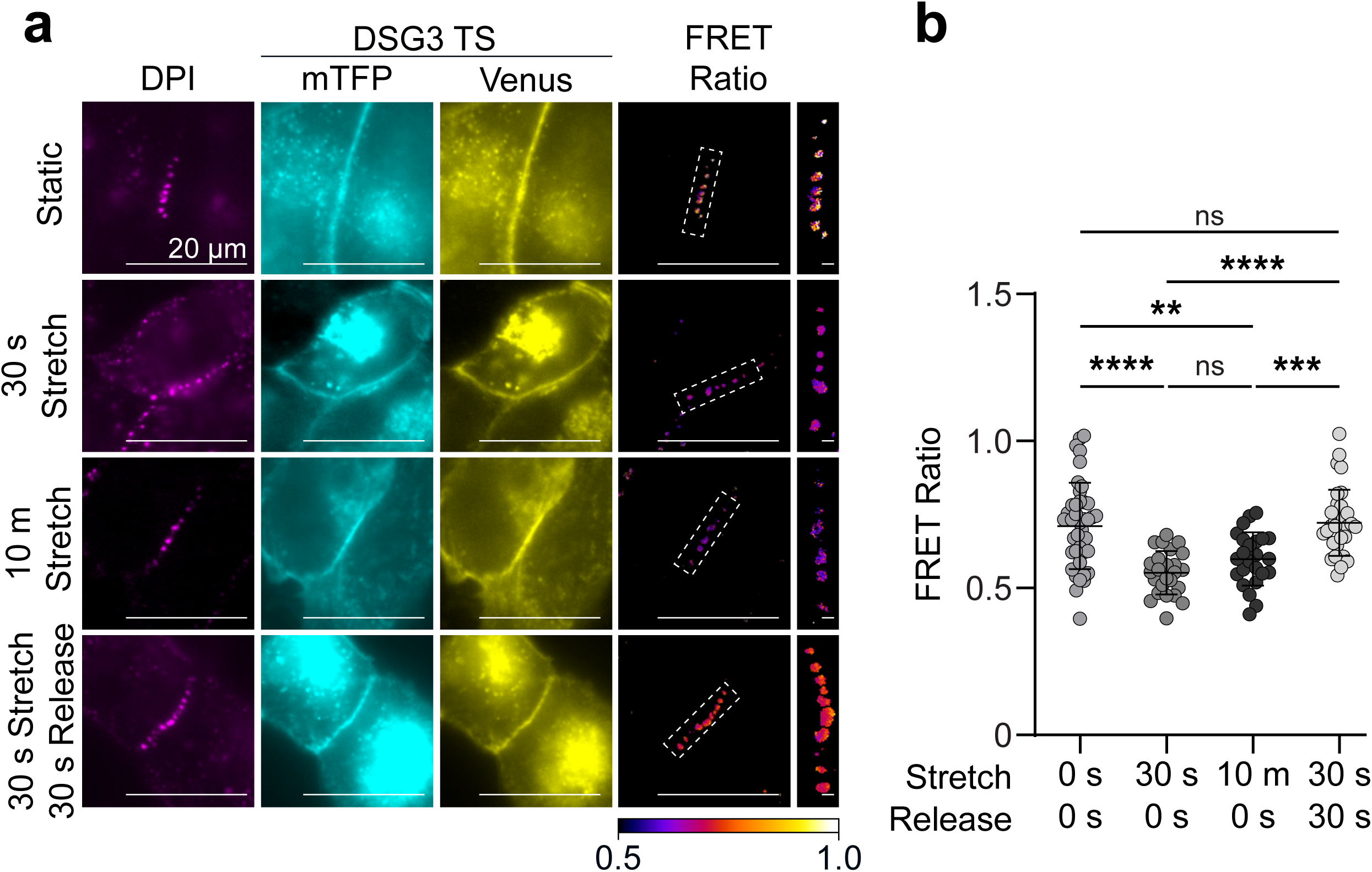
The opening of the DP head domain leads to tension on Dsg3. **a**, Representative images of DPI-Halo cells expressing DSG3-TS constructs in static, 30 s stretch no release, 10 m stretched no release, and 30 s stretch with 30 s released junctions. Scale bar = 20 μm for full image, scale bar = 1 μm for FRET ratio ROI. **b.** Scatter plot of the FRET ratio per image of DSG3-TS expressed in DPI-Halo cells (Static: n = 41 0.71 ± 0.15; 30 s stretch no release: n = 25, 0.55 ± 0.07; 10 m stretched no release: n = 24 0.6 ± 0.09; 30 s stretch with 30 s Release: n = 33 0.72 ± 0.11. n = FRET ratio; mean ± sd). Shapiro-Wilks test confirmed normal distribution (p<0.05). A Brown-Forsythe and Welch ANOVA with Dunnett’s T3 multiple comparisons test assessed significant differences in FRET ratio (**** p<0.0001, *** p<0.001, ** p<0.01, ns not significant,)

**Figure 7:**
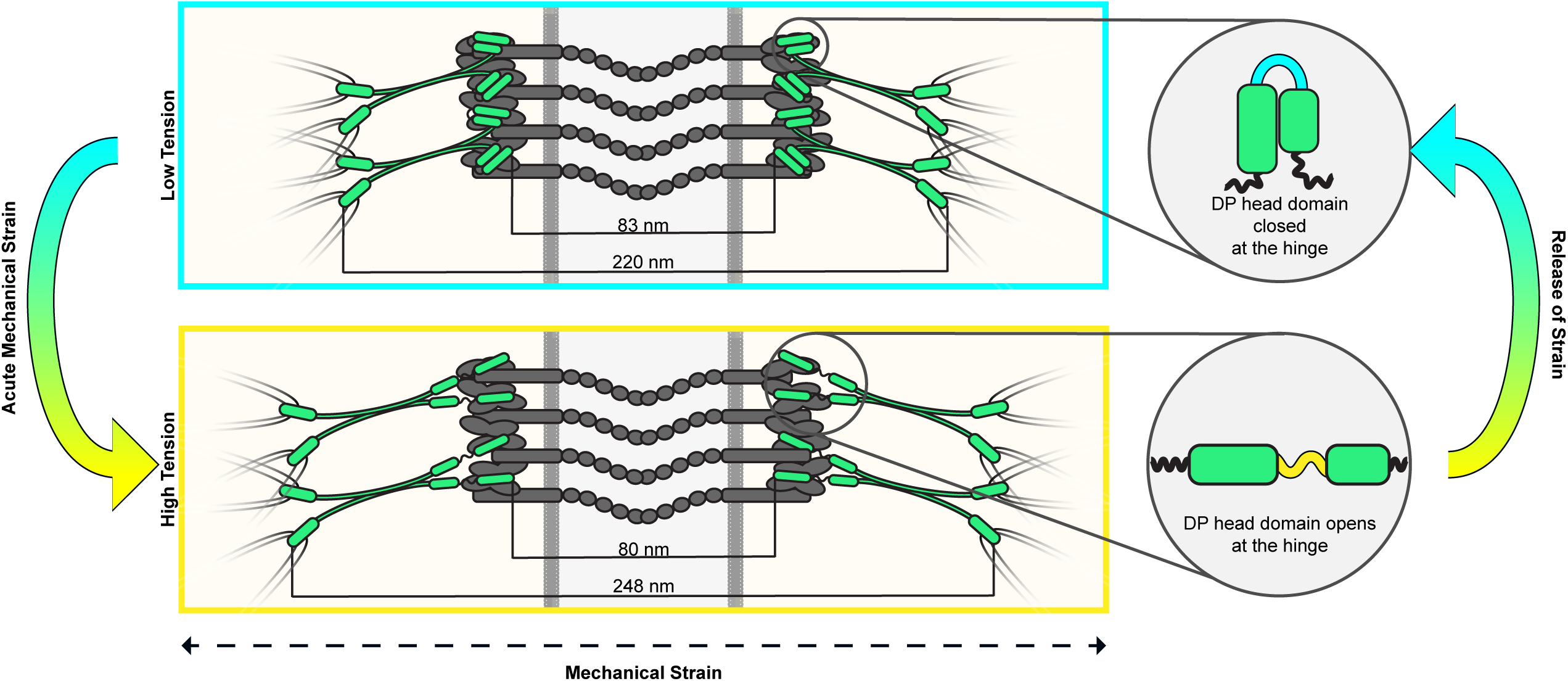
The cycle of desmosomal force response. Diagram describing how desmosomes respond and reset to induction and release of acute mechanical forces. Note how after experiencing acute mechanical force, the DP head domain opens leading to a repositioning of the tail domain. Following release of force, the head domain closes leading to the tail domain position returning back to baseline.

The proposed opening and closing of the head domain is supported by *in vitro* work identifying folded “U” and extended “I” conformations from purified DP head domain^10^ and MD simulations, which revealed an opening of the DP head domain in response to mechanical stretch^7^. Here we used novel DP variants to show the hinge is responsible for tail domain repositioning in response to mechanical forces. Interestingly, the purified envoplakin head domain adopts only one conformation that reflects a tight right angle between its long and short arms^10^. Here we show that substituting the DP hinge with the envoplakin hinge abrogated DP’s mechanical response, strongly supporting the key role of the full-length DP hinge in desmosome function.

We found that while the DP tail domain position can change during maturation, the hinge-dependent repositioning of the DP tail observed in this study is a specific response to acute mechanical force. The maturation-induced phenotype was preserved with all DP variants tests, indicating tail domain repositioning during maturation is controlled by a separate mechanism, perhaps ODP binding proteins leading to changes in the orientation of the head domain, which is read out as the tail domain movements during maturation.

Mechanosensitivity is an important regulatory feature in other adhesive cell junctions. Alpha catenin (a-cat) and talin link adherens junctions and focal adhesions, respectively, to F-actin and undergo mechanical opening under strain^19–27^. The MI domain of a-cat is opened by mechanical forces, exposing a cryptic vinculin binding domain leading to vinculin recruitment and stabilization of adhesion^19,20^. The rod domains of talin are mechanically sensitive with 13 helical bundles that undergo reversible unfolding upon mechanical force^25^. Once unfolded, vinculin binding domains are exposed that allow for talin stabilization^23,24^. Talin opening has an additional consequence of repositioning of its tail domain, with going from an all-closed state (50 nm) to all-open state (350 nm), resulting in a change in distance from the plasma membrane^22^. This change in positioning is proposed to play a role in cell signaling. Talin and a-cat mechanosensitivity are important for adhesion and reinforcing stability of their adhesion complexes^21,23,27^.

Two non-exclusive hypotheses can explain the role of the DP head domain opening in demosomes: one is mechanical, the other biochemical. We found the head domain opens and closes rapidly in response to acute mechanical forces. First, this opening will change the way forces are transmitted through the desmosome. The physical movement of the tail domain away from the membrane in response to force may relieve tension across the entire junction. Additionally, the head domain opening will ultimately change the impulse of an acute mechanical force on other protein-protein interactions, spreading the acceleration of force over time. This would dampen the forces cascading through the desmosome from keratin to DP to the cadherins, ultimately protecting desmosomal cadherin *trans*-binding.

Second, the DP head domain opening and closing may expose cryptic binding sites or targets for post translational modifications. Binding with known ODP binding partners, plakophilins, plakoglobin, and the desmosomal cadherin tails could be strengthened, or currently unidentified proteins may be recruited. This could serve to reinforce the ODP and protect the junction against mechanical forces. Cryptic binding sites on either arm of the DP head domain or within the hinge could be revealed by strain. Desmosomes have been linked to YAP and therefore the Hippo pathway, with Dsg 3 forming a complex with pYAP^28^, Additionally, DP binds with the guanine exchange factor Ect2, promoting active RhoA proximal to desmosomes^18^. YAP acts as a mechanosensor for focal adhesions and adherens junctions, making it an attractive candidate for regulation via DP mechanoresponse^29^.

Skin can experience multiple types of forces: acute loading applied suddenly as investigated here, sustained loading where the tissue is held in a stretched position over a period of time, and cyclic loading where the tissue is repeatedly stretched and released. These forces are also multi-dimensional and often irregular. Reversibility of desmosome architecture would be critical to “reset” desmosomes after one force to be ready for whatever forces come next. We also found the DP architectural change is dependent on desmosome orientation relative to the stretch axis. This is similar to established changes in the magnitude of DP tension relative to mechanical force^30^. This represents an agile spatially localized mechanism for combating mechanical forces, capable of quick resting and response to many sources.

This work provides novel insights into desmosome mechanotransduction, identifying the DP hinge as critical for the reversible repositioning of its tail domain under force. DP head domain opening provides mechanical strength to cells and may serve as an initial buffer for force propagation throughout the complex. Integration of how desmosomal proteins respond to mechanical forces remains a critical barrier to our understanding of dynamic cell adhesion. Our discovery of how mechanical forces impact DP architecture provides a new framework for the molecular mechanisms underlying the ability of desmosomes to provide mechanical integrity to cells and tissues.

## Methods

### Cell Culture

HaCaT cells were cultured in Dulbecco’s modified Eagle’s medium (DMEM; Corning, Tewksbury, MA, USA) supplemented with 10% fetal bovine serum (FBS; A5670801, Gibco) and 2% Penicillin/streptomycin (15070063, Gibco) at 37°C and 5% CO2. DPI-mEGFP constructs have a C-terminal mEGFP following a linker (DPPVAT) in the pREP[EXP]-CMV mammalian gene expression vector from Vector Builder (Chicago, IL, USA). DPI-Halo was generated from DPI-mEGFP by inserting a halo tag followed by a stop codon following the DPPV in the flexible linker. Stable cells were generated by transfecting the constructs into DP KO HaCaT cells using Lipofectamine 3000 (LC3000015; Thermo Fisher Scientific, Waltham, MA, USA). For FRET experiments, DPI-Halo stably expressing cells were transfected with a Dsg3 tension sensor construct using Lipofectamine 3000 according to the manufacturer’s protocols. To enrich the GFP-positive or Halo tag positive populations, cells were sorted two days following the initial transfection with fluorescence activated cell sorting (FACS, BD FacsAria III; BD BioSciences, Franklin Lakes, NJ, USA). To maintain expression of the DPI constructs, cells were cultured in DMEM supplemented with 500 ug/ml G418 (10131027, Gibco) and 10% FBS.

### Construct Generation

Halo-tagged WT DPI-mEGFP construct were generated using Thermo Scientific™ Phusion™ Hot Start II DNA Polymerase (F549L, Thermo Fisher Scientific, Waltham, MA, USA). The HaloTag sequence was amplified from pHalo2X-N3 (251420, Addgene, Watertown, MA, USA) and cloned into the WT DPI-mEGFP using AgeI restriction enzyme, with a stop codon introduced at the 3′ end of the HaloTag sequence preventing mEGFP expression.

### Cell Stretching

Cells were seeded onto PDMS membranes (SC-0022, STREX USA, San Diego, CA, USA) coated with fibronectin and cultured for 2 days to achieve confluence. The membranes were placed on sterilized glass coverslips and affixed to the cell stretching rig (ST-1500, STREX USA, San Diego, CA, USA). The membranes underwent a 30% uniaxial stretch at 1000 μm/s. Cells were fixed at either 30 s or 5 m in the stretched position or released after 30 s of stretch for 30 s or 5 m. Following fixation, membranes were removed from the cell stretching system and processed for imaging.

### Antibodies

Antibodies for immunofluorescence: FluoTag-X4 anti-GFP conjugated with Alexa Fluor 647 (used for dSTORM, 1:250, N0304 NanoTag Biotechnologies, Gottingen, Germany), FluoTag-X4 anti-GFP conjugated with Atto488 (1:250, N0304 NanoTag Biotechnologies, Gottingen, Germany), anti-desmoplakin (1:150, A303-356A Bethyl Laboratories, Montgomery, TX, USA), anti-desmoplakin (N-terminus, 1:400, 73403 Cell Signaling, Danvers, MA, USA), Anti-DSG3 (1:400, 20483 Cell Signaling, Danvers, MA, USA), anti-rabbit Alexa Fluor 647 (1:1000; a-21244 Thermo Fisher Scientific, Waltham, MA, USA), anti-mouse Alexa Fluor 647 (1:1000; a-21235 Thermo Fisher Scientific, Waltham, MA, USA)

### Immunofluorescence

Cells were cultured on either glass coverslips or PDMS membranes coated with fibronectin. At either 1, 2, or 5 days of growth in a confluent culture, samples were fixed with either 4% paraformaldehyde (PFA; Electron Microscopy Sciences, Hatfield, PA, USA) or a 1:1 acetone: methanol cocktail for 10 m. All samples grown on PDMS membranes were fixed at room temperature as well as samples grown on glass fixed with 4% PFA. Samples fixed with 1:1 acetone: methanol were fixed at 4°C. Following fixation, samples were washed 3x (5 m) at RT with washing buffer (PBS supplemented with 0.5% triton X-100, 0.5% normal goat serum (NGS; 16210-064, Gibco), and 0.5% normal horse serum (NHS; 16050-130, Gibco)). Samples then incubated in blocking buffer (PBS supplemented with 0.5% triton X-100, 5% normal goat serum, 5% normal horse serum, and 10 mg/ml BSA) for 30 m at RT. For primary antibody labeling, samples were incubated in either humidity chambers at 37°C for 2 h with rocking or overnight at 4°C based upon antibody recommendations. Following primary antibody labeling, samples were washed 3x (5 m) at RT with washing buffer. For secondary antibody labeling, samples were incubated with secondary antibody diluted in blocking buffer for 1 h at 37°C. Samples were washed 3x with PBS and stored at 4°C protected from light until imaging.

### Halo Tag Treatment

DPI-Halo stable cells were washed with Hank’s Balanced Salt Solution (HBSS; 14-170-161, Thermo Fisher Scientific, Waltham, MA, USA). Samples then incubated in DMEM supplemented with 500 ug/ml G418, 10% FBS, and 50 mM Janelia Fluor 635 HaloTag Ligand (HT1050; Promega, Madison, WI, USA) for 1 h 37°C and 5% CO2.

### Microscopy

Confocal images were acquired on a Nikon Ti-2 AXR microscope with a 60x 1.42 NA oil immersion objective, with 405, 488, and 647 nm laser excitation with Nyquist sampling. Z-stacks were acquired and deconvolved with the Richardson-Lucy algorithm with eight iterations.

dSTORM images were obtained on a Nikon Ti-2 microscope system with a 100x 1.49 NA oil immersion objection, 647 nm laser, and Andor iXon EMCCD camera. For each image, 10,000 frames were acquired. Samples were imaged in a 50 mM Tris-HCl pH 8.0, 10 mM NaCl, 10% glucose buffer with 5% 1 M MEA and 2% GLOX (20% 17 mg/ml catalase and 14 mg glucose oxidase), each prepared in a 50 mM Tris-HCl and 10 mM NaCl buffer.

FRET experiments were obtained on a Nikon Ti-2 microscope system with a 60x 1.42 NA oil immersion objective, with 405, 488, and 647 nm laser excitation. For each image group, an image was captured containing donor only excitation and emission, donor excitation and acceptor emission, acceptor excitation and emission, and Janelia Fluor 635 excitation and emission.

### FRET Analysis

For each image, a 15 px rolling ball background subtraction was applied. A binary mask was generated from the DPI-Halo. The FRET ratio image was calculated by dividing the donor excitation-acceptor emission image by the donor excitation-donor emission image. The DP mask was applied to the FRET ratio image, and the average FRET ratio across all regions in each image was measured.

### Confocal Image Analysis

15-slice maximum projections of each condition were generated within ImageJ. The distance around cells with desmosomes with multiple cell-cell borders were measured and the number of junctions for each cell was counted.

### dSTORM Image Analysis

dSTORM data were reconstructed at 4 nm/pixel and analyzed using our in-house MATLAB software. Individual desmosomes were manually selected, and a rectangular ROI was drawn to enclose both plaques and align with the desmosome midline. Fluorescence intensity was averaged parallel to the midline to generate an intensity profile perpendicular to the plaques. Profiles were smoothed, and plaque-to-plaque distance was measured as the separation between the two intensity peaks flanking the ROI center, converted to nm using the reconstruction pixel size.

### Dispase Fragmentation Assay

Cells were seeded, grown to confluency, and treated with 2.4 U/ml dispase (D4693 Millapore Sigma, St. Louis, MO, USA) until cell sheets are lifted. Using an automated p1000 pipette (Eppendorf, Hamberg, Germany) with a FBS coated tip, cell sheets were fragmented. Samples are then fixed with 1% PFA, stained with 0.2% Methylene blue overnight. Fragments were counted and imaged the following day.

### Statistical Analysis

GraphPad Prism was used for all statistical analysis. Figure legends report all statistical tests used, mean ± sd for each dataset, *n*, data point representation, and significance analysis (**** p<0.0001, *** p<0.001, ** p<0.01, *p<0.05, ns – not significant). Shapiro-Wilks test for normality confirmed if data were normally distributed. For datasets that contained non-normally distributed samples, Mann-Whitney and Kruskal-Wallis with Dunn’s multiple comparison tests were performed. Welch’s t-test or One-way ANOVA with Dunnett’s multiple comparisons test was conducted on normally distributed datasets.

## Data and Resource availability

All relevant data is found within the article and supplementary material

## Acknowledgements

The authors thank the High-Resolution Imaging Facility at the University of Alabama at Birmingham for their support in dSTORM imaging. The UAB High Resolution Imaging Facility was supported by NCI P30 CA013148.

## Contributions

Conceptualization: C.M.A., A.L.M.;

Formal analysis: C.M.A., W.F.D.;

Funding acquisition: A.L.M., C.M.A.;

Investigation: C.M.A., Y.T.B.T., S.C.B.;

Methodology: C.M.A., W.F.D., E.J.L., A.L.M., D.E.C., A.P.K.;

Resources: D.E.C.;

Supervision: A.L.M.;

Validation: C.M.A., Y.T.B.T., S.C.B.;

Visualization: C.M.A., W.F.D., S.C.B., A.L.M.;

Writing – original draft: C.M.A.;

Writing – review & editing: C.M.A., W.F.D., Y.T.B.T., S.C.B., E.J.L., A.P.K., D.E.C., A.L.M.

---

**Fig S1.**
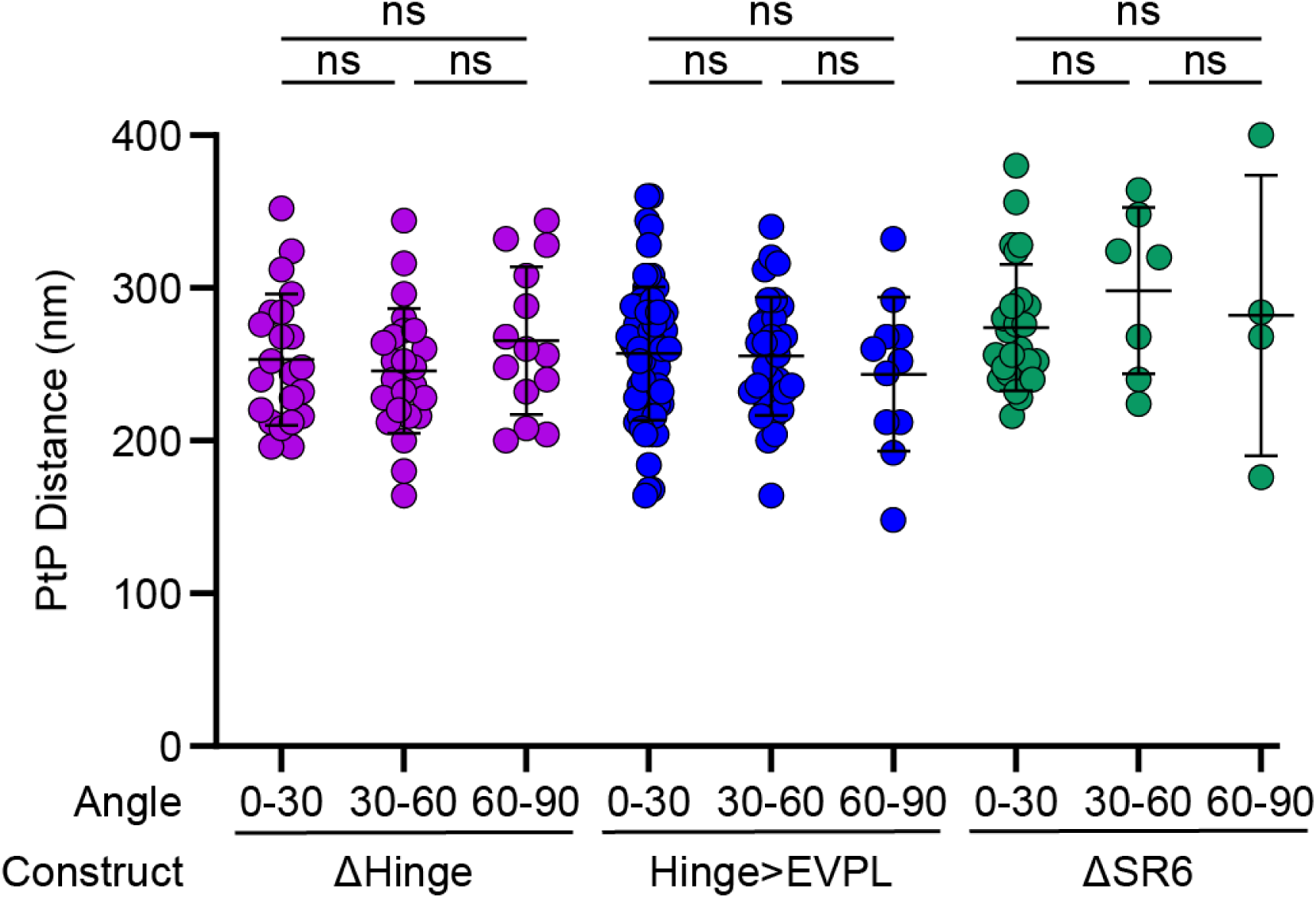
DP Head Domain Constructs Sorted by Angle Relative to Stretch. Scatter plot of the tail domain plaque-to-plaque (PtP) distances of stretched DPI-mEGFP head domain constructs sorted by angle relative to uniaxial stretch (ΔHinge stretched: 0-30: n = 22, 253 ± 43 nm; ΔHinge stretched 30-60: n = 24, 248 ± 41 nm; ΔHinge stretched 60-90: n = 14, 265 ± 49 nm; Hinge>EVPL stretched 0-30: n = 66, 257 ± 44 nm; Hinge>EVPL stretched 30-60: n = 32, 255 ± 39 nm; Hinge>EVPL stretched 60-90: n = 11, 244 ± 50 nm; ΔSR6 stretched 0-30: n = 25, 274 ± 41nm; ΔSR6 stretched 30-60: n = 7, 298 ± 54 nm; ΔSR6 stretched 60-90: n = 4, 282 ± 92 nm. n is number of desmosomes, mean ± sd). Shapiro -Wilks test assessed normality. One-way ANOVA with Tukey’s multiple comparison test revealed there was no statistical differences in PtP distances of Hinge>EVPL and ΔHinge when sorted by orientation to stretch axis (ns not significant). Kruskal-Wallis with Dunn’s multiple comparison test revealed no statistical difference PtP distance of ΔSR6 when sorted by orientation to stretch axis (ns not significant).

**Fig S2.**
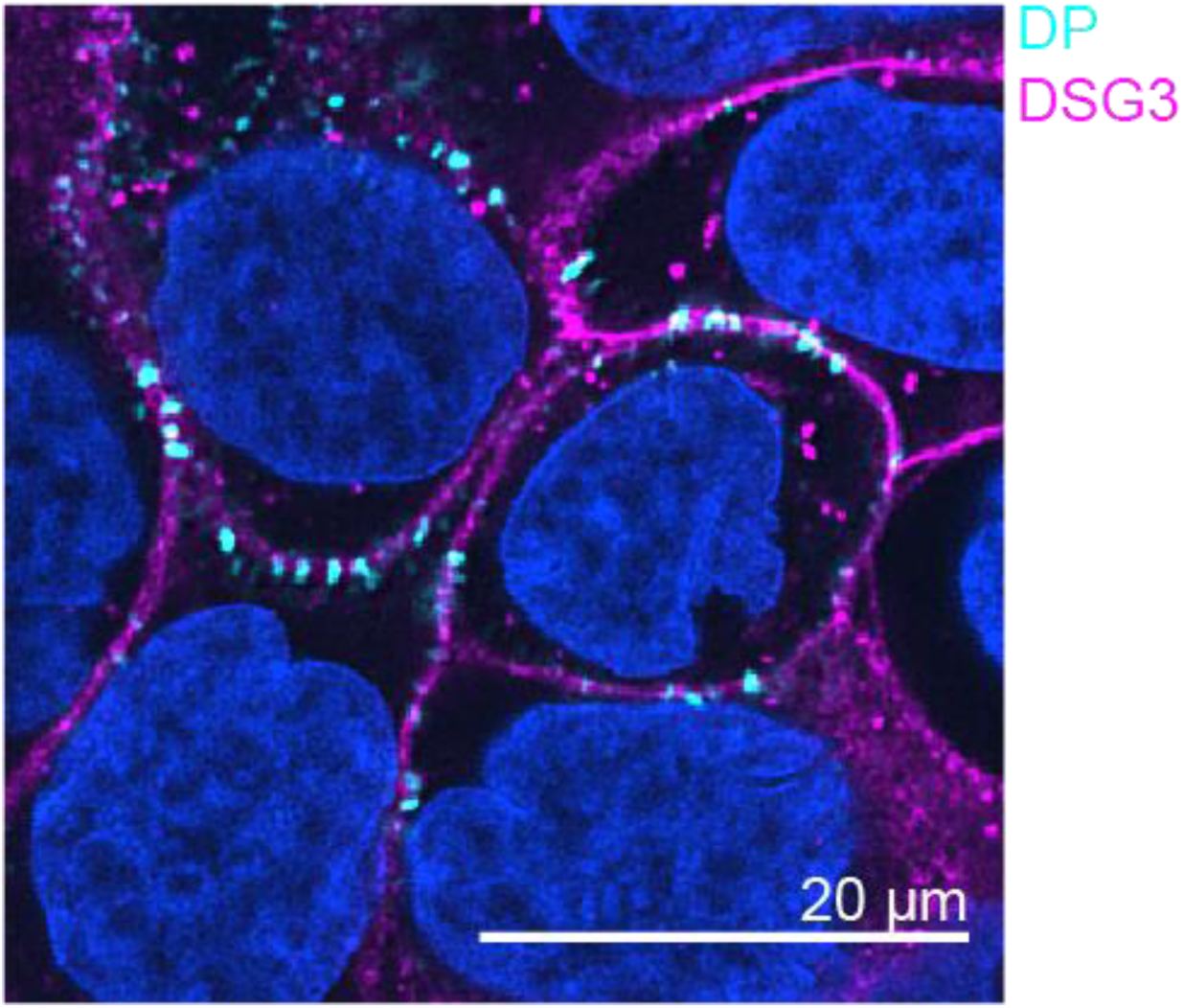
Representative confocal max projection of DPI-Halo stable cells with Dsg 3(DP-cyan, Dsg3-magenta). Scale bar = 20 µm

**Fig. S3.**
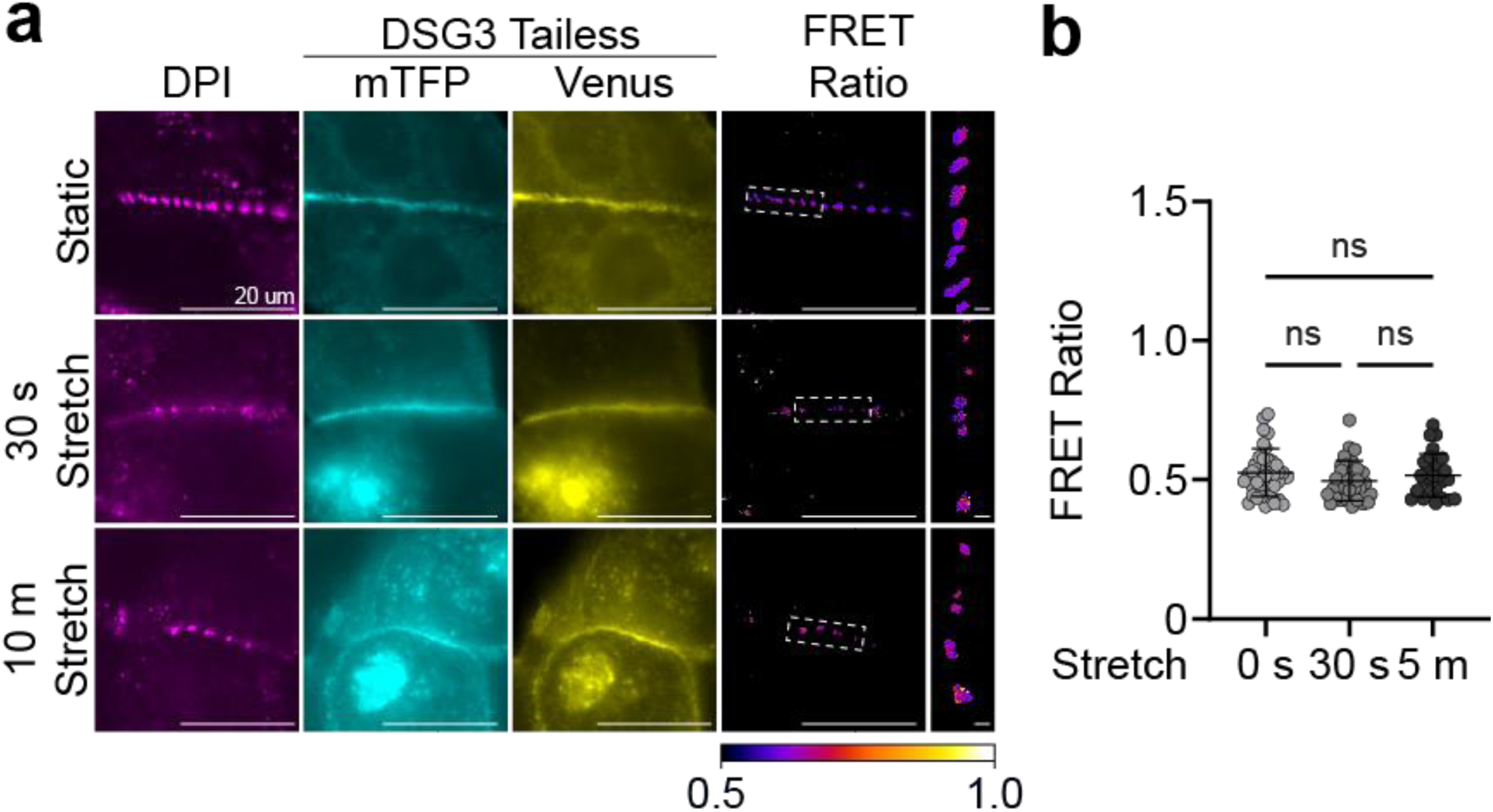
Dsg3-TL Tension does not Change to Mechanical Strain. **a,** Representative images of DPI-Halo cells expressing Dsg 3-TL constructs in static, 30 s stretch, and 10 m stretch junctions. Scale bar = 20 µm, ROI scale bar = 1 µm **b,** Scatter plot of the FRET ratio per image of Dsg 3-TL expressed in DPI-Halo Cells (static, = 40, 0.5 ± 0.1; 30 s stretch n = 31, 0.5 ± 0.1; 10 m stretch, n = 27, 0.5 ± 0.1. n = image analyzed, mean ± sd). Shapiro-wilks test assessed normality. Kruskal-Wallis with Dunn’s multiple comparison test revealed no statistical difference in FRET ratio between conditions (ns = not significant).

## References

1. Roberts N, Horsley V. Developing stratified epithelia: lessons from the epidermis and thymus. Wiley Interdiscip Rev Dev Biol. 2014;3(6):389–402. Epub 20140829. doi: 10.1002/wdev.146. PubMed PMID: 25176390; PMCID: PMC4283209.

2. Kowalczyk AP, Green KJ. Structure, function, and regulation of desmosomes. Prog Mol Biol Transl Sci. 2013;116:95–118. doi: 10.1016/B978-0-12-394311-8.00005-4. PubMed PMID: 23481192; PMCID: PMC4336551.

3. Stahley SN, Kowalczyk AP. Desmosomes in acquired disease. Cell Tissue Res. 2015;360(3):439–56. Epub 20150321. doi: 10.1007/s00441-015-2155-2. PubMed PMID: 25795143; PMCID: PMC4456195.

4. Broussard JA, Getsios S, Green KJ. Desmosome regulation and signaling in disease. Cell Tissue Res. 2015;360(3):501–12. Epub 20150219. doi: 10.1007/s00441-015-2136-5. PubMed PMID: 25693896; PMCID: PMC4489137.

5. Baddam SR, Arsenovic PT, Narayanan V, Duggan NR, Mayer CR, Newman ST, Abutaleb DA, Mohan A, Kowalczyk AP, Conway DE. The Desmosomal Cadherin Desmoglein-2 Experiences Mechanical Tension as Demonstrated by a FRET-Based Tension Biosensor Expressed in Living Cells. Cells. 2018;7(7). Epub 20180626. doi: 10.3390/cells7070066. PubMed PMID: 29949915; PMCID: PMC6070948.

6. Jin X, Rosenbohm J, Kim E, Safa BT, Moghaddam AO, Seiffert-Sinha K, Leiker M, Jones E, Zhai H, Baddam SR, Minnick G, Huo Y, Wahl JK, 3rd, Meng F, Huang C, Lim JY, Conway DE, Sinha AA, Yang R. Desmosomal cadherin tension loss in pemphigus vulgaris mediated by the inhibition of active RhoA at cell-cell adhesions. iScience. 2025;28(8):113081. Epub 20250709. doi: 10.1016/j.isci.2025.113081. PubMed PMID: 40740498; PMCID: PMC12307676.

7. Dong Y, Elgerbi A, Xie B, Han Y, Kwiatkowski AV, Choy JS, Sivasankar S. Actomyosin forces trigger a conformational change in desmoplakin within desmosomes. Nat Commun. 2025;16(1):9052. Epub 20251010. doi: 10.1038/s41467-025-64124-4. PubMed PMID: 41073558; PMCID: PMC12514153.

8. Seeley LD, Ainslie CM, Sewell-Loftin MK, Mattheyses AL. Super-resolution imaging reveals stretch-induced architectural rearrangement of desmoplakin in desmosomes. J Invest Dermatol. 2026. Epub 20260430. doi: 10.1016/j.jid.2026.04.016. PubMed PMID: 42069078.

9. Green KJ, Stappenbeck TS, Parry DA, Virata ML. Structure of desmoplakin and its association with intermediate filaments. J Dermatol. 1992;19(11):765–9. doi: 10.1111/j.1346-8138.1992.tb03777.x. PubMed PMID: 1293163.

10. Al-Jassar C, Bernadomicron P, Chidgey M, Overduin M. Hinged plakin domains provide specialized degrees of articulation in envoplakin, periplakin and desmoplakin. PLoS One. 2013;8(7):e69767. Epub 20130729. doi: 10.1371/journal.pone.0069767. PubMed PMID: 23922795; PMCID: PMC3726778.

11. Kang H, Weiss TM, Bang I, Weis WI, Choi HJ. Structure of the Intermediate Filament-Binding Region of Desmoplakin. PLoS One. 2016;11(1):e0147641. Epub 20160125. doi: 10.1371/journal.pone.0147641. PubMed PMID: 26808545; PMCID: PMC4726743.

12. Al-Jassar C, Bikker H, Overduin M, Chidgey M. Mechanistic basis of desmosome-targeted diseases. J Mol Biol. 2013;425(21):4006–22. Epub 20130802. doi: 10.1016/j.jmb.2013.07.035. PubMed PMID: 23911551; PMCID: PMC3807649.

13. Mohammed F, Chidgey M. Desmosomal protein structure and function and the impact of disease-causing mutations. J Struct Biol. 2021;213(3):107749. Epub 20210524. doi: 10.1016/j.jsb.2021.107749. PubMed PMID: 34033898.

14. Bharathan NK, Mattheyses AL, Kowalczyk AP. The desmosome comes into focus. J Cell Biol. 2024;223(9). Epub 20240809. doi: 10.1083/jcb.202404120. PubMed PMID: 39120608; PMCID: PMC11317759.

15. North AJ, Bardsley WG, Hyam J, Bornslaeger EA, Cordingley HC, Trinnaman B, Hatzfeld M, Green KJ, Magee AI, Garrod DR. Molecular map of the desmosomal plaque. J Cell Sci. 1999;112 (Pt 23):4325–36. doi: 10.1242/jcs.112.23.4325. PubMed PMID: 10564650.

16. Beggs RR, Rao TC, Dean WF, Kowalczyk AP, Mattheyses AL. Desmosomes undergo dynamic architectural changes during assembly and maturation. Tissue Barriers. 2022;10(4):2017225. Epub 20220105. doi: 10.1080/21688370.2021.2017225. PubMed PMID: 34983311; PMCID: PMC9621066.

17. Stahley SN, Bartle EI, Atkinson CE, Kowalczyk AP, Mattheyses AL. Molecular organization of the desmosome as revealed by direct stochastic optical reconstruction microscopy. J Cell Sci. 2016;129(15):2897–904. Epub 20160617. doi: 10.1242/jcs.185785. PubMed PMID: 27505428; PMCID: PMC5004873.

18. Zarkoob H, Kam CY, Koetsier JL, McCarthy E, Jaiganesh A, Kelsell DP, Sheikh F, Godsel LM, Green KJ. RhoGEF Ect2 supports RhoA activity at cell-cell junctions through desmoplakin. Life Sci Alliance. 2026;9(6). Epub 20260312. doi: 10.26508/lsa.202503454. PubMed PMID: 41819613; PMCID: PMC12983042.

19. Yao M, Qiu W, Liu R, Efremov AK, Cong P, Seddiki R, Payre M, Lim CT, Ladoux B, Mege RM, Yan J. Corrigendum: Force-dependent conformational switch of alpha-catenin controls vinculin binding. Nat Commun. 2015;6:6568. Epub 20150304. doi: 10.1038/ncomms7568. PubMed PMID: 25735633.

20. Pang SM, Le S, Kwiatkowski AV, Yan J. Mechanical stability of alphaT-catenin and its activation by force for vinculin binding. Mol Biol Cell. 2019;30(16):1930–7. Epub 20190718. doi: 10.1091/mbc.E19-02-0102. PubMed PMID: 31318313; PMCID: PMC6727763.

21. del Rio A, Perez-Jimenez R, Liu R, Roca-Cusachs P, Fernandez JM, Sheetz MP. Stretching single talin rod molecules activates vinculin binding. Science. 2009;323(5914):638–41. doi: 10.1126/science.1162912. PubMed PMID: 19179532; PMCID: PMC9339221.

22. Margadant F, Chew LL, Hu X, Yu H, Bate N, Zhang X, Sheetz M. Mechanotransduction in vivo by repeated talin stretch-relaxation events depends upon vinculin. PLoS Biol. 2011;9(12):e1001223. Epub 20111220. doi: 10.1371/journal.pbio.1001223. PubMed PMID: 22205879; PMCID: PMC3243729.

23. Hirata H, Tatsumi H, Lim CT, Sokabe M. Force-dependent vinculin binding to talin in live cells: a crucial step in anchoring the actin cytoskeleton to focal adhesions. Am J Physiol Cell Physiol. 2014;306(6):C607–20. Epub 20140122. doi: 10.1152/ajpcell.00122.2013. PubMed PMID: 24452377.

24. Atherton P, Stutchbury B, Wang DY, Jethwa D, Tsang R, Meiler-Rodriguez E, Wang P, Bate N, Zent R, Barsukov IL, Goult BT, Critchley DR, Ballestrem C. Vinculin controls talin engagement with the actomyosin machinery. Nat Commun. 2015;6:10038. Epub 20151204. doi: 10.1038/ncomms10038. PubMed PMID: 26634421; PMCID: PMC4686655.

25. Yao M, Goult BT, Klapholz B, Hu X, Toseland CP, Guo Y, Cong P, Sheetz MP, Yan J. The mechanical response of talin. Nat Commun. 2016;7:11966. Epub 20160707. doi: 10.1038/ncomms11966. PubMed PMID: 27384267; PMCID: PMC4941051.

26. Ishiyama N, Sarpal R, Wood MN, Barrick SK, Nishikawa T, Hayashi H, Kobb AB, Flozak AS, Yemelyanov A, Fernandez-Gonzalez R, Yonemura S, Leckband DE, Gottardi CJ, Tepass U, Ikura M. Force-dependent allostery of the alpha-catenin actin-binding domain controls adherens junction dynamics and functions. Nat Commun. 2018;9(1):5121. Epub 20181130. doi: 10.1038/s41467-018-07481-7. PubMed PMID: 30504777; PMCID: PMC6269467.

27. Duong CN, Bruckner R, Schmitt M, Nottebaum AF, Braun LJ, Meyer Zu Brickwedde M, Ipe U, Vom Bruch H, Scholer HR, Trapani G, Trappmann B, Ebrahimkutty MP, Huveneers S, de Rooij J, Ishiyama N, Ikura M, Vestweber D. Force-induced changes of alpha-catenin conformation stabilize vascular junctions independently of vinculin. J Cell Sci. 2021;134(24). Epub 20211222. doi: 10.1242/jcs.259012. PubMed PMID: 34851405; PMCID: PMC8729784.

28. Uttagomol J, Ahmad US, Rehman A, Huang Y, Laly AC, Kang A, Soetaert J, Chance R, Teh MT, Connelly JT, Wan H. Evidence for the Desmosomal Cadherin Desmoglein-3 in Regulating YAP and Phospho-YAP in Keratinocyte Responses to Mechanical Forces. Int J Mol Sci. 2019;20(24). Epub 20191210. doi: 10.3390/ijms20246221. PubMed PMID: 31835537; PMCID: PMC6940936.

29. Nardone G, Oliver-De La Cruz J, Vrbsky J, Martini C, Pribyl J, Skladal P, Pesl M, Caluori G, Pagliari S, Martino F, Maceckova Z, Hajduch M, Sanz-Garcia A, Pugno NM, Stokin GB, Forte G. YAP regulates cell mechanics by controlling focal adhesion assembly. Nat Commun. 2017;8:15321. Epub 20170515. doi: 10.1038/ncomms15321. PubMed PMID: 28504269; PMCID: PMC5440673.

30. Price AJ, Cost AL, Ungewiss H, Waschke J, Dunn AR, Grashoff C. Mechanical loading of desmosomes depends on the magnitude and orientation of external stress. Nat Commun. 2018;9(1):5284. Epub 20181211. doi: 10.1038/s41467-018-07523-0. PubMed PMID: 30538252; PMCID: PMC6290003.

